# A targeted e-learning approach to reduce student mixing during a pandemic

**DOI:** 10.1101/2020.06.10.135533

**Authors:** Sing Chen Yeo, Clin K.Y. Lai, Jacinda Tan, Joshua J. Gooley

## Abstract

The COVID-19 pandemic has resulted in widespread closure of schools and universities. These institutions have turned to distance learning to provide educational continuity. Schools now face the challenge of how to reopen safely and resume in-class learning. However, there is little empirical evidence to guide decision-makers on how this can be achieved. Here, we show that selectively deploying e-learning for larger classes is highly effective at decreasing campus-wide opportunities for student-to-student contact, while allowing most in-class learning to continue uninterrupted. We conducted a natural experiment at a large university that implemented a series of e-learning interventions during the COVID-19 outbreak. Analyses of >24 million student connections to the university Wi-Fi network revealed that population size can be manipulated by e-learning in a targeted manner according to class size characteristics. Student mixing showed accelerated growth with population size according to a power law distribution. Therefore, a small e-learning dependent decrease in population size resulted in a large reduction in student clustering behaviour. Our results show that e-learning interventions can decrease potential for disease transmission while minimizing disruption to university operations. Universities should consider targeted e-learning a viable strategy for providing educational continuity during early or late stages of a disease outbreak.

## Main

The coronavirus disease 2019 (COVID-19) has had enormous socioeconomic impact^1^. In 5 months, 5.5 million COVID-19 cases have been confirmed, resulting in 350,000 deaths across 188 countries^2^. The severe acute respiratory syndrome coronavirus 2 (SARS-CoV-2) that causes COVID-19 is primarily spread when an infected person sneezes or coughs^3^. The spread of infection can be slowed by public health measures that reduce person-to-person contact. Nonpharmaceutical interventions that include restricted travel, staying at home, and physical distancing can delay and flatten the peak of COVID-19 cases to avoid the overwhelming of medical services^4–6^. Nonpharmaceutical interventions therefore play a critical role in controlling the spread of disease until effective vaccines or drugs are available^7^.

School closure is a key strategy for controlling the spread of infectious diseases^7–9^. Empirical and modelling studies show that closing schools and universities can suppress COVID-19 transmission when combined with other nonpharmaceutical interventions^5,6,10^. The COVID-19 pandemic has resulted in an unprecedented number of school and university closures, affecting over 1.2 billion learners worldwide^11^. Consequently, a massive shift from classroom learning to distance learning has occurred^12^. This has created great strain on educational institutions which function not only as places of learning, but also as major employers and drivers of local economies. It is therefore important to consider less disruptive interventions to ensure educational continuity^13^. However, there is a major knowledge gap in how student mixing patterns on campus are affected by school policies enacted during the disease outbreak. Evidence-based solutions are needed for schools and universities to re-open safely as soon as possible.

We conducted a natural experiment to evaluate the impact of implementing e-learning measures on student population dynamics at the National University of Singapore (NUS) during the COVID-19 outbreak. NUS is the largest university in Singapore, with about 24,000 undergraduate students per year enrolled in course modules with in-class learning (**Extended Data Table 1**). In line with the national public health response, NUS adopted nonpharmaceutical interventions that aimed to reduce risk of SARS-CoV-2 transmission (**Extended Data Table 2**). Normal in-class learning took place during the first 4 weeks of the semester, coinciding with the first imported case of COVID-19 (**Fig. 1a**). Shortly afterward the first local transmission of COVID-19 was identified. This escalated the nationwide pandemic response and prompted NUS to implement e-learning over the next several weeks for all classes with >50 students. As the number of COVID-19 cases continued to climb in Singapore and globally as a pandemic, NUS implemented e-learning for all classes with >25 students. One week later, nationwide ‘enhanced circuit-breaker’ measures were announced, which led to the suspension of all in-class learning.

**Fig. 1.**
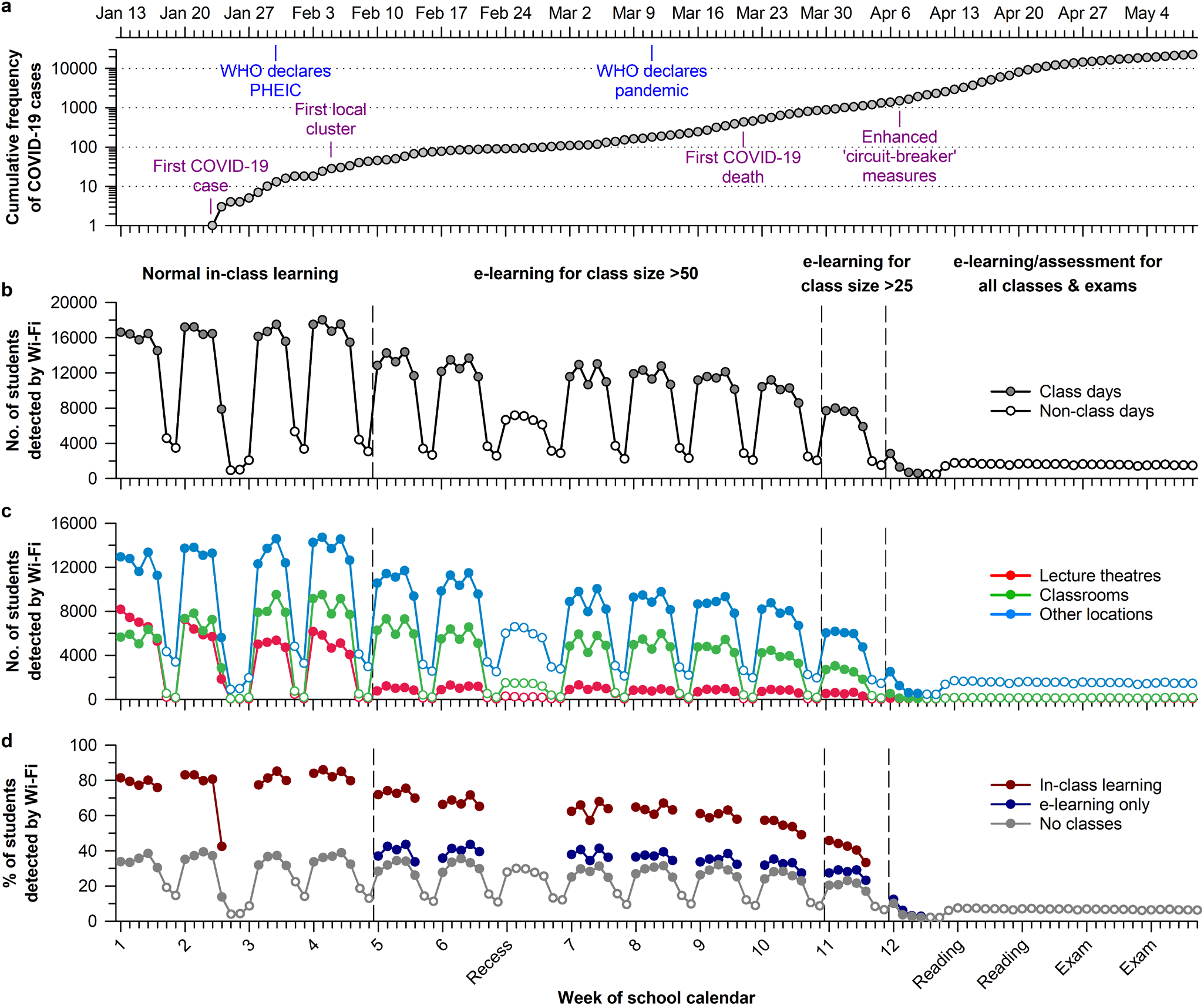
E-learning interventions decreased the number of students detected on campus during the COVID-19 outbreak. (**a**) The timeline of COVID-19 cases in Singapore is shown for the second semester of the 2019/20 school year at the National University of Singapore (NUS). Each e-learning intervention was associated with a decrease in the daily number of students who connected to the NUS Wi-Fi network, assessed (**b**) campus-wide and (**c**) for different types of locations on campus. (**d**) The daily percentage of students detected by Wi-Fi was about two-fold greater in students with at least one class conducted by in-class learning, as compared with students with e-learning only or no scheduled class. In panels **c** and **d**, open circles indicate non-class days. COVID-19, Coronavirus Disease 2019; WHO, World Health Organisation; PHEIC, Public Health Emergency of International Concern.

The multi-phased transition to e-learning during the COVID-19 outbreak provided a unique opportunity to investigate student mixing patterns on campus. Social interactions and their products show super-linear growth with population size^14–18^. Hence, interventions that reduce the number of students by a small amount would be expected to decrease substantially the potential for person-to-person contact and disease transmission. We predicted that the shift to e-learning would result in a moderate drop in the number of students on campus due to fewer students having to attend in-class sessions. We further predicted that this would result in a large drop in opportunities for student mixing. These predictions were tested by analysing >24 million student connections to the NUS Wi-Fi network, comprising several thousand Wi-Fi access points across campus. Each time that a student connected to the Wi-Fi network, the time and location data were recorded. These data were used to investigate students’ spatiotemporal mixing patterns before and during the disease outbreak.

In the early part of the semester, there were about 16,500 students per school day who connected to the NUS Wi-Fi network (**Fig. 1b**). The only notable exception was the eve of the Chinese New Year holiday, in which the number of students detected by Wi-Fi dropped by about half. After the transition to e-learning for classes with >50 students, there was a 30% decrease in the number of students detected on campus. Wi-Fi connections decreased sharply in lecture theatres and moderately in classrooms and non-teaching facilities (**Fig. 1c**). The number of students detected by Wi-Fi dropped by an additional 25% after e-learning was implemented for classes with >25 students. These findings contrast with results from the previous academic year in which the daily number of students detected by Wi-Fi on school days was stable across the semester (**Extended Data Fig. 1**).

Effects of e-learning on the number of students on campus were determined by students’ class sizes and schedules. The transition to e-learning for classes with >50 or >25 students impacted a small proportion of total classes (9% and 19%, respectively) but these classes had high student enrolment (**Extended Data Fig. 2**). Therefore, most students had at least one class that was converted to e-learning. On a typical school day, nearly 18,000 students had a scheduled class compared with 6,000 students with no class (**Extended Data Fig. 2**). Due to heterogeneity in students’ timetables, the transition to e-learning resulted in a subset of students each day with classes delivered only by e-learning (**Fig. 1d**). These students were detected on campus at about the same rate as those who had no class (about 35-40%). In contrast, students with in-class learning were detected at nearly twice the rate as those students with e-learning only or no class (about 60-80%). Linear regression analysis showed that 90% of the variance in the daily number of students detected on campus was explained by the number of students with in-class learning (**Extended Data Fig. 3**).

Next, we evaluated the impact of e-learning on student clustering behaviour. A student cluster was defined as >25 students connected to the same Wi-Fi access point. We surveyed several thousand Wi-Fi access points to determine the number of sites with student clustering and the duration of clustering at each of these sites (**Extended Data Fig. 4**). There were several hundred Wi-Fi access points where clustering occurred, with 20% of these sites accounting for about 80% of the total duration of clustering behaviour over the semester (**Extended Data Fig. 5**). The daily rhythm in number of students on campus drove the pattern of clustering behaviour (**Fig. 2a**). In the early part of the semester, student clustering tracked the timing of lectures, whereas this pattern was flattened with e-learning (**Fig 2b**).

**Fig. 2.**
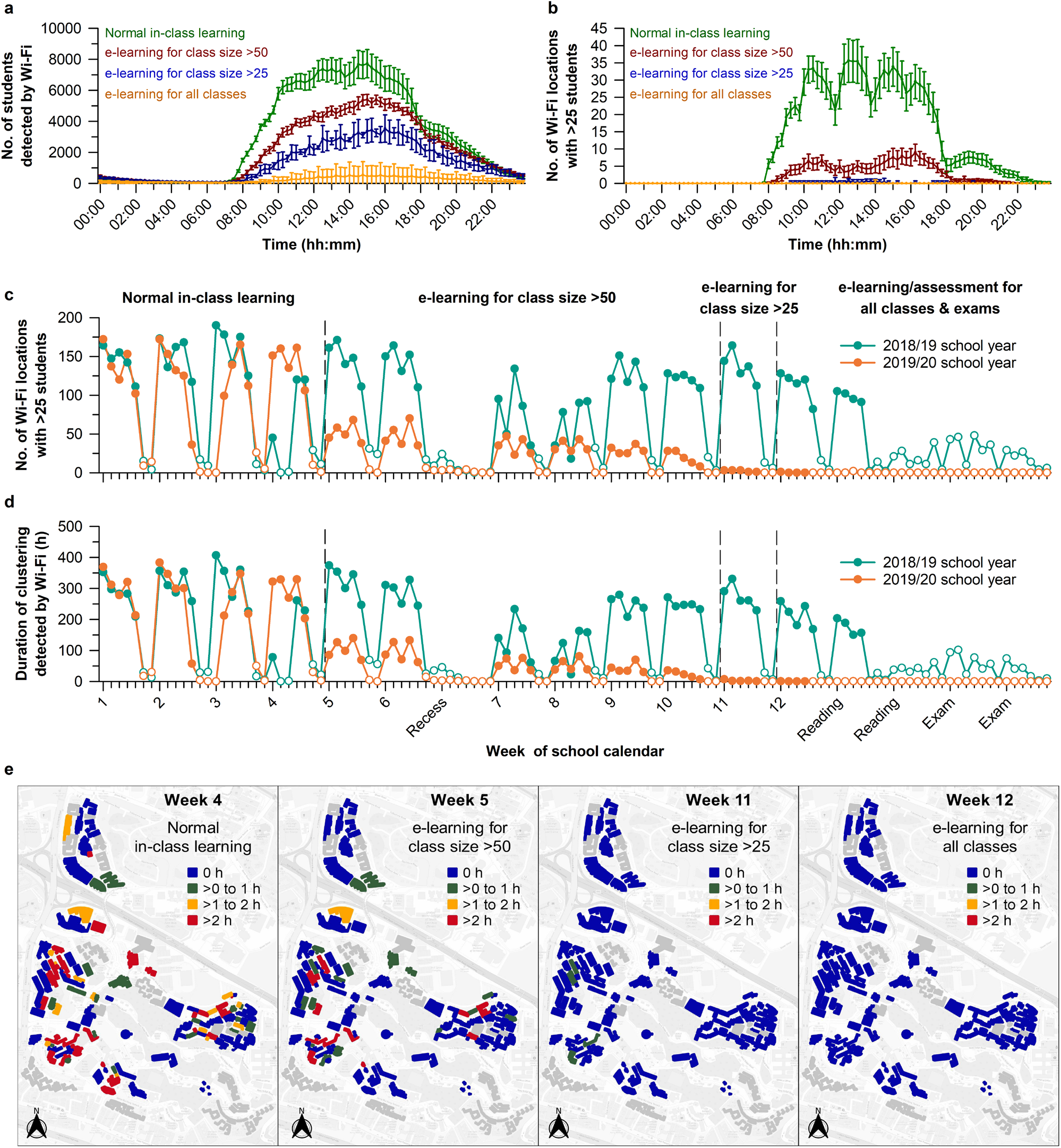
E-learning interventions reduced student clustering during the COVID-19 outbreak. Each e-learning intervention was associated with (**a**) a decrease in the daily rhythm in students detected on campus, and (**b**) a flattening in the daily time course of locations with >25 students connected to the same Wi-Fi access point. E-learning measures during the disease outbreak (2019/20 school year) were effective at decreasing (**c**) the daily number of Wi-Fi locations with a student cluster, and (**d**) the duration of clustering at these sites, as compared with the prior academic year with normal in-class learning (2018/19 school year). (**e**) The daily duration of student clustering in campus buildings decreased as more stringent e-learning policies were implemented. In panels **a** and **b**, the daily mean ± 95% CI is shown for different parts of the semester. In panels **c** and **d**, open circles indicate non-class days. In panel **e**, buildings are color-coded by the daily average of clustering duration. Buildings with missing or incomplete Wi-Fi data are coloured grey.

During normal in-class learning, there were about 150 Wi-Fi access points per day where a student cluster was detected (**Fig. 2c**), contributing to about 300 hours of clustering behaviour (**Fig. 2d**). The transition to e-learning for classes with >50 students was associated with a 70% decrease in the number of sites with a student cluster, as well as the duration of clustering at these sites. These findings differ from the prior academic year, in which student clustering behaviour on school days changed little over the semester (**Fig. 2c-d**). The transition to e-learning for classes with >25 students effectively eliminated student clustering. These findings were further visualised by plotting the data on university map to identify hot spots of clustering activity (**Fig. 2e**). After e-learning, there was a marked reduction in student clustering in buildings where students usually converged for classes and social activities.

Each e-learning transition was associated with a decrease in the number of unique pairs of students with spatiotemporal overlap (**Fig. 3a**). Over a typical day, nearly half a million unique pairs of students showed Wi-Fi connection overlap. This number was cut in half after e-learning was implemented for classes with >50 students, and it dropped further as more restrictive e-learning policies were enacted. We then examined the degree of overlap for individual students (i.e., the number of unique pairs formed by a student), focusing on the top 100 students per day with the greatest amount of spatiotemporal overlap with their peers (**Fig. 3b**). Each e-learning transition was associated with a decrease in the degree of student overlap, with network plots demonstrating weakening of the spatiotemporal student network (**Fig. 3c**).

**Fig. 3.**
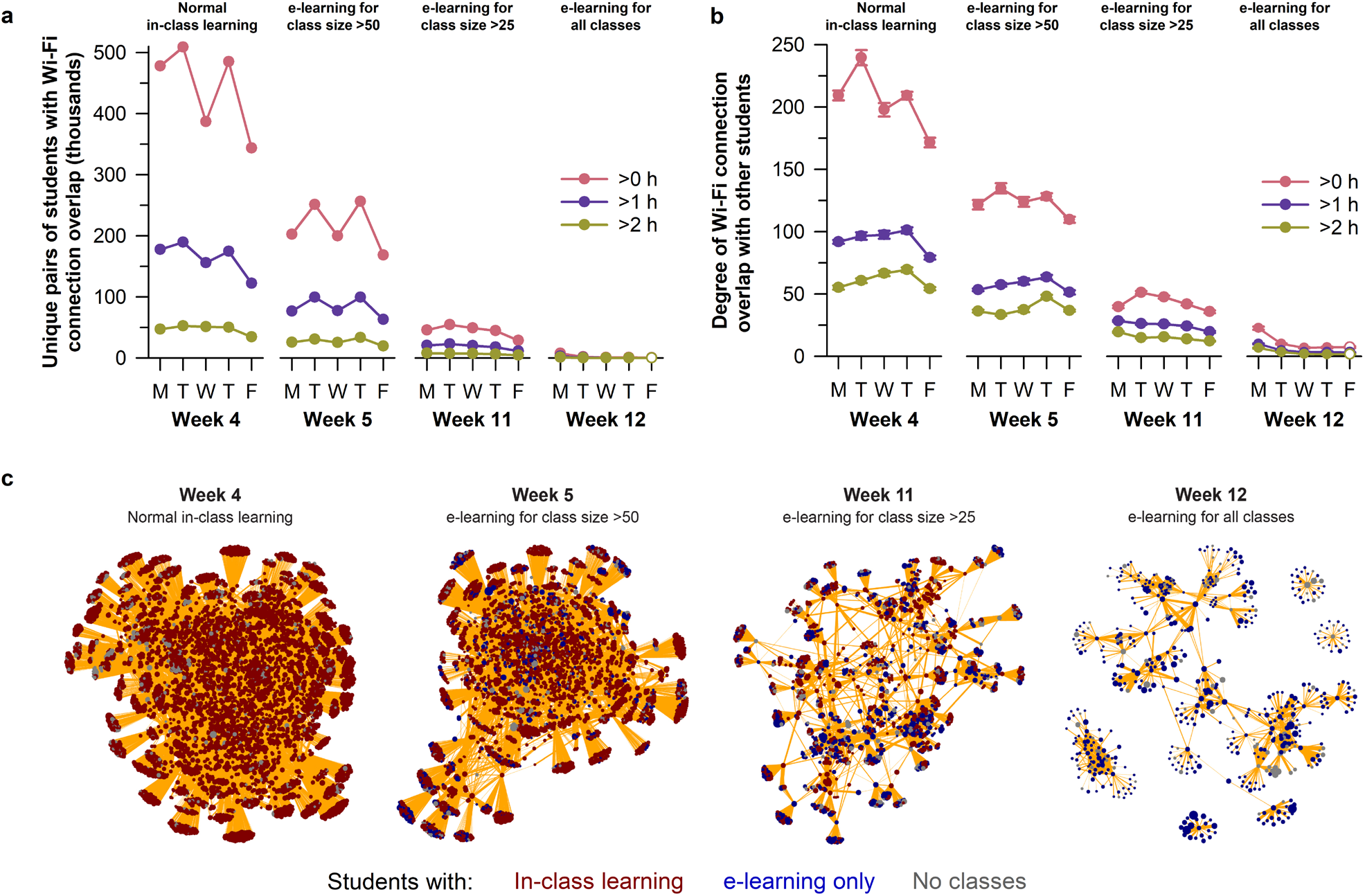
E-learning interventions reduced the number of pairs of students with spatiotemporal overlap during the COVID-19 outbreak. (**a**) Each e-learning intervention was associated with a decrease in the daily number of unique pairs of students who connected simultaneously to the same Wi-Fi access point. In the top 100 students per day with the greatest degree of Wi-Fi connection overlap with their peers, e-learning interventions were associated with (**b**) a decrease in the number of students with whom they shared overlap, and (**c**) a sparser student network structure. In panels **a** and **b**, results are plotted for different durations of Wi-Fi connection overlap between students. In panel **b**, the mean ± 95% CI is shown for each group of 100 students. In panel **c**, the size of each circle relates to the daily duration of time connected to Wi-Fi. The thickness of the orange lines corresponds to the duration of spatiotemporal overlap between pairs of students. Data in the network plots correspond to Mondays for each of the representative weeks with different e-learning policies.

Next, we investigated scaling properties of student mixing patterns with the number of students detected on campus. The number of Wi-Fi access points with student clustering increased with student population size according to a power law distribution (**Fig. 4a**). The relationship was super-linear whereby growth in the number of student clusters accelerated with larger numbers of students on campus. Similar results were observed for the daily duration of student clustering (**Fig. 4b**). These findings were reproducible using data from the prior academic year, demonstrating that scaling properties of student clustering behaviour with population size were generalisable and not related to the COVID-19 pandemic or implementation of e-learning (**Extended Data Fig. 6**). Power law scaling of student clustering behaviour was also observed for different types of locations on campus including teaching and non-teaching facilities (**Extended Data Fig. 7**).

**Fig. 4.**
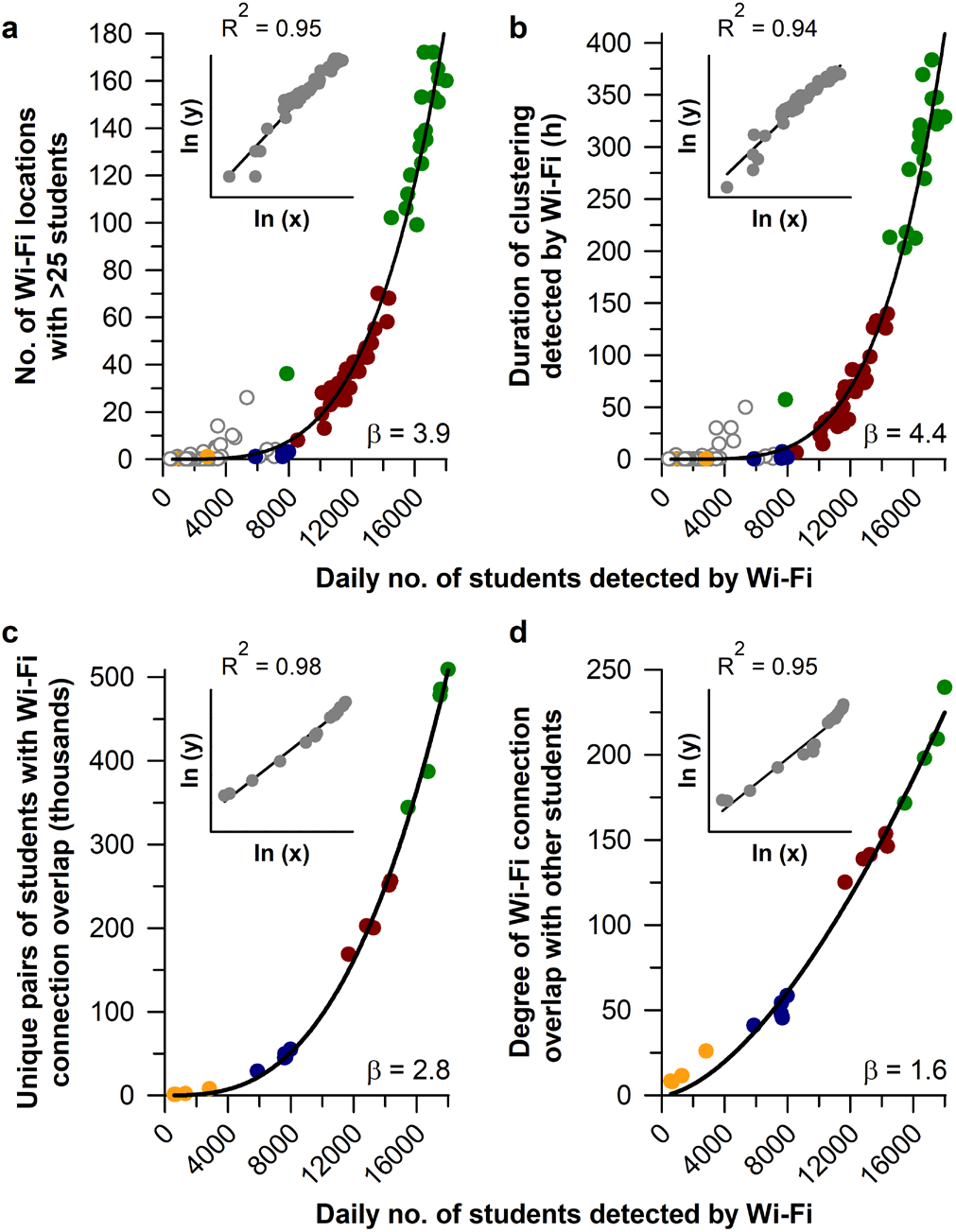
Student clustering showed power law scaling with the number of students on campus. Student mixing showed accelerated growth with daily population size, including (**a**) the number of Wi-Fi locations with a student cluster, (**b**) the duration of student clustering, (**c**) the number of unique pairs of students who connected to the same Wi-Fi access point, and (**d**) the degree of spatiotemporal overlap among the top 100 students per day who overlapped most with their peers. Data are shown for the second semester of the 2019/20 school year during the COVID-19 outbreak. Each dataset was fitted with a power law function, with β representing the scaling exponent. Insets show results for linear regression after taking the natural logarithm of each variable. Circle colours correspond to different parts of the semester with normal in-class learning (green), e-learning for classes with >50 students (red), e-learning for classes with >25 students (blue), and e-learning for all classes (orange). In panels **a** and **b**, open circles indicate non-class days.

Moreover, power law scaling was observed when alternative definitions of cluster size were tested, ranging from >5 to >50 students detected at the same Wi-Fi access point (**Extended Data Fig. 8**). These analyses showed that larger clusters of students were more sensitive to changes in population size (i.e., the exponent of the power law function was greater), and that e-learning resulted in a marked decrease in the frequency and duration of clustering behaviour for all cluster sizes. In line with these observations, the number of unique pairs of students with spatiotemporal overlap exhibited super-linear scaling with daily population size (**Fig. 4c**), as did the degree of overlap for ‘highly-connected’ individual students with their peers (**Fig. 4d**).

The scaling properties of student mixing patterns have important implications for strategies that seek to minimize person-to-person contact during a disease outbreak. We showed that a small decrease in student population size resulted in a large reduction in student clustering behaviour. Hence, an important goal for reducing risk of disease transmission is to decrease the number of students on campus. This can be achieved in a predictable manner by implementing e-learning for all classes that exceed a given class size. It is more practical to focus on larger classes because they are often conducted in a lecture format that can be converted easily to e-learning (e.g., video lecture), and they are a main driver of student clustering behaviour on campus.

The power law scaling we observed is consistent with prior work demonstrating accelerated growth of human interactions with city population size^14–18^. Epidemiological models indicate that these scaling relationships drive super-linear growth of disease transmission rates as cities get bigger^15,17,18^. Like cities, universities are complex systems composed of different infrastructural and social elements whose hierarchical structures give rise to scaling laws^14,19^. However, we found that the growth rates of student mixing patterns on campus (determined by the exponent of the power law scaling function) were greater compared with studies on scaling of human interactions with city size. This may be related to differences in student network dynamics and university infrastructural components compared with cities in which they reside. Earlier work examined social connectivity patterns derived from mobile phone call records and internet interactions^16–18^. Such methods capture information about social networks but not the potential for physical contact between individuals. By comparison, Wi-Fi connection data provide information on student proximity patterns, which is more relevant for assessing disease transmission risk.

Our study is the first to measure campus-wide spatiotemporal mixing of students using the university’s Wi-Fi network. At its core, this method requires counting of students connected to different Wi-Fi access points. Students are only detected if they have a Wi-Fi enabled device that is actively scanning for a Wi-Fi access point. The location where a student is connected also depends on the proximity and range of the nearest Wi-Fi access point. Despite these limitations, prior studies have shown that Wi-Fi connections are as accurate as dedicated physical sensors (e.g., infrared beam-break or thermal sensors) for estimating student occupancy of university rooms and buildings^20,21^. The daily pattern of student Wi-Fi connections also conforms to expectations for different sites on campus including teaching spaces, libraries, food courts, and residential buildings^20,22–24^. An important limitation of our study is that we did not investigate student mixing with university staff or visitors because we only had Wi-Fi data for students. In future work, it will be important to evaluate the scaling properties of clustering behaviour while considering all persons on campus. Here, we took advantage of the university’s existing Wi-Fi network infrastructure to collect data from students without the need for their active participation. This approach can be adopted for continual monitoring of students’ Wi-Fi connection patterns and clustering behaviour, and it can be extended to include all other users of the Wi-Fi network.

In conclusion, e-learning is an intervention that universities can use to provide educational continuity while decreasing student-to-student contact during a disease outbreak. We recommend that e-learning be incorporated into each university’s pandemic preparedness plan. First, universities should evaluate their class size distribution to determine the impact of a given e-learning policy on the daily number of students with in-class learning. This information makes it possible to achieve a targeted reduction in student population size because the number of students on campus is dependent on the proportion of students with in-class learning. Second, universities should develop the capacity to count the number of students on campus because population size is a main driver of student clustering behaviour and mixing patterns. This can be achieved using existing Wi-Fi network infrastructure, and the data can be used to derive scaling properties of student mixing with population size. Third, universities should periodically perform e-learning exercises during normal operations. This can be used to measure the impact of e-learning on student mixing, and to test infrastructure required for campus-wide e-learning during an epidemic^25^. Fourth, universities should consider how to implement e-learning in view of the local and nationwide health response to a disease outbreak. A partial transition to e-learning may be most appropriate before widespread community spread has occurred, or during recovery to normal school operations. A full transition to e-learning may be required near the peak of an epidemic. Taken together, our study establishes a roadmap that universities can follow for making evidence-based decisions on students’ learning and safety during the COVID-19 pandemic and future disease outbreaks.

## Methods

### Student data and ethics statement

Our study was performed using university archived data managed by the NUS Institute for Applied Learning Sciences and Educational Technology (ALSET). The ALSET Data Lake stores and links deidentified student data across different university units for the purpose of conducting educational analytics research^26^. Data tables in the ALSET Data Lake are anonymised by student tokens which map identifiable data to a hash string using a one-way function that does not allow recovery of the original data. The same student-specific tokens are represented across tables, allowing different types of data to be combined without knowing students’ identities. The data types used in our study included basic demographic information (age, sex, ethnicity, citizenship, year of matriculation), class enrolment information, and Wi-Fi connection metadata. Students included in our study provided informed consent to the NUS Student Data Protection Policy, which explains that student data may be used for research and evaluating university policies. Research analyses were approved by the NUS Learning Analytics Committee on Ethics.

### Timeline of COVID-19 cases and university policies

The timeline of COVID-19 cases in Singapore was determined using daily situation reports published online by the Ministry of Health (MOH)^27,28^. Nationwide alerts and policies regarding the public health response were taken from press releases available on the MOH website^29^. University policies enacted during the COVID-19 outbreak were compiled from circulars distributed to staff and students, and they are archived by the NUS Office of Safety, Health, and Environment^30^.

### Student timetables and class size characteristics

Student data were analysed in the second semester of the 2018/19 and 2019/20 school years. This allowed us to compare student behaviour before and during the COVID-19 outbreak over an equivalent period (from January to May). Students’ class schedules and class sizes were derived from student enrolment data provided by the NUS Registrar’s Office. At NUS, students enrol in course modules, many of which are further divided into different lectures, class groups, tutorials, or laboratory sessions. We analysed data in students taking at least one module that required in-class learning (23,668 and 23,993 students in 2018/19 and 2019/20 school years). Data were excluded from students taking only fieldwork or project-based modules with no in-class component (2,722 and 3,240 students). Class size was defined as the number of students who were scheduled to meet in the same place for a given course module. The timing and location of classes were retrieved using the NUSMods application programming interface (https://api.nusmods.com/v2/). Timetable data were sorted for each school day of the semester to identify students with scheduled in-class learning. These data were also used to determine which classes were converted to e-learning based on class size. This allowed us to calculate the daily number of students with in-class learning, e-learning only, or no class.

### Wi-Fi connection data

Connections to the NUS Wi-Fi network are continually monitored by NUS Information Technology to evaluate and improve services provided to the university. The campus-wide wireless network comprises several thousand Wi-Fi access points and deploys different types of routers (*Cisco* Aironet 1142, 2702I and 2802I) and wireless protocols (802.11n 2.4 GHz, 802.11n 5 GHz, and 802.11ac 5 GHz). Each time that a person’s Wi-Fi enabled device associates with the NUS wireless network the transmission data are logged. Students’ Wi-Fi connection metadata were added daily to the ALSET Data Lake by a data pipeline managed by NUS Information Technology. Each data point included the tokenized student identity, the anonymized media access control (MAC) address used to identify the Wi-Fi enabled device of the student (e.g., smartphone, tablet, or laptop), the name and location descriptor of the Wi-Fi access point, and the start and end time of each Wi-Fi connection. The name and location descriptor usually carried information about the room or building in which the Wi-Fi access point was located. By cross-referencing these data with the known timing and location of classes, we categorised Wi-Fi access points into teaching facilities (lecture theatres or classrooms) and non-teaching facilities.

### Analyses of student mixing patterns

The Wi-Fi dataset comprised more than 24 million student connections to the wireless network over 2 semesters. Students’ connection data were binned in 15-min intervals to reduce the size of the data, resulting in 11,328 epochs that spanned 118 days in each semester. In instances where students were connected to more than one Wi-Fi access point in the same epoch, they were assigned to the access point in which their Wi-Fi enabled device received the greatest volume of data (i.e., based on megabytes of data received). The resulting table of Wi-Fi connections and access points was used to derive time and location information for each student over the semester. This enabled us to count the daily number of students who connected to the Wi-Fi network, and the number of students who were connected to the same Wi-Fi access point within a 15-min epoch. The latter was used to examine student clustering behaviour. We defined a cluster as >25 students connected to the same Wi-Fi access point because of the high potential for student-to-student contact, and it aligned with the university’s e-learning policy prior to suspension of in-class learning (i.e., e-learning for class size >25). The duration of student clustering at each Wi-Fi access point was calculated as the sum of 15-min epochs with >25 students. Data were analysed using R statistical software (version 3.6.3)^31^.

Geospatial clustering was visualised by plotting students’ data on a map of the NUS campus. The researchers did not have access to the geospatial coordinates for Wi-Fi access points. Therefore, general location information provided in the Wi-Fi metadata (e.g., name of the building or room) was used to determine manually the building locations. Using sources that included the official NUS campus map and venues listed on class timetables, we confirmed the geospatial coordinates for 80% of Wi-Fi access points. Georeferencing was performed by mapping Wi-Fi access points to vector point shapefiles representing individual buildings. The ESRI shapefiles required for mapping were obtained from the OpenStreetMap geodatabase for the region of Malaysia, Singapore, and Brunei (map tiles in the OpenStreetMap are licensed under CC BY-SA www.openstreetmap.org/copyright, © OpenStreetMap contributor). We used QGIS software (version 3.12.1) to edit the vector points and to insert names of Wi-Fi access points to the attribute table. Student clustering within each building was determined by pooling the duration of clustering across all Wi-Fi access points within the building. Subsequently, we merged the clustering duration data with the ESRI shapefiles using the “sf” package (version 0.9-0)^32^ in the R software environment. The QGIS platform was then used to visualise student clustering for 124 buildings across the NUS campus. Buildings with incomplete Wi-Fi data and student hostels were excluded from the analysis.

The number of unique pairs of students with spatiotemporal overlap in their Wi-Fi connections was determined for 4 representative weeks of the semester (weeks 4, 5, 11, 12). These time intervals captured the transition from normal in-class learning to e-learning for classes with >50 students (week 4 to 5), and the transition from e-learning for classes with >25 students to e-learning for all classes (week 11 to 12). The decision to focus on these temporal windows was driven by practical reasons related to computing resources required to analyse the data. In each student, the degree of Wi-Fi connection overlap was determined by counting the number of unique students with whom he/she shared a Wi-Fi connection. Our analyses focused on the top 100 students per day with the greatest degree of overlap with their peers because we expected this group would illustrate best the impact of e-learning on individual student networks. This student group size was also practical for visualising effects of e-learning on student network structure, which was performed using the “igraph” package^33^ (version 1.2.5) with the force-directed layout algorithm (layout_with_fr) in the R software environment.

Student clustering behaviour on school days was modelled as a function of daily student population size using a power law scaling equation: *y* = *aN*^*β*^. In this equation, *y* is the measure of student mixing (e.g., number of Wi-Fi access points with a student cluster, duration of student clustering, or pairs of students with Wi-Fi connection overlap); *a* is a constant; *N* is the daily population size estimated by the number of students who connected to the NUS Wi-Fi network; and the exponent *β* reflects the underlying dynamics (e.g., hierarchical structure, social networks, and infrastructure) of the university ecosystem. We considered other mathematical functions, including exponential and hyperbolic equations, but they did not fit as well to the data. Variables that show power law scaling are linearly related when each variable is logarithmically transformed. We therefore took the natural logarithm of each pair of variables (i.e., the student mixing variable and daily population size) and performed linear regression to confirm the expected linear relationship. The coefficient of determination (R^2^ value) was used to evaluate goodness-of-fit for the regression model. Modelling and regression analyses were performed using Sigmaplot software (Version 14; Systat Software, Inc) and R statistical software.

## Data availability

The data that support the findings of this study will be made available from the corresponding author upon reasonable request. Requests will be handled in compliance with data sharing and data management policies of the National University of Singapore.

## Acknowledgements

The work was supported by funding provided by the NUS Office of the Senior Deputy President & Provost and the NUS Institute for Applied Learning Sciences and Educational Technology (ALSET). We thank research and administrative staff at ALSET and NUS Information Technology (NUS IT). We thank Kevin Hartman (ALSET) and Yung Shing Gene (NUS IT) for assisting with data management and troubleshooting.

## Author contributions

S.C.Y. and C.K.L analysed the data. S.C.Y, C.K.L, and J.T. prepared tables, figures, and other source materials required for the paper. J.J.G. designed the research and wrote the paper. All authors read and approved the final manuscript.

---

**Extended Data Table 1.** Student characteristics and general information.

|  | 2018/19 school year<br>(*n* = 23,668) | 2019/20 school year<br>(*n* = 23,993) |
| --- | --- | --- |
| **Demographic information** |  |  |
| Age in years (mean ± SD) | 22.7 ± 2.1 | 21.6 ± 1.9 |
| Sex, *n* (%) |  |  |
| Female | 12,179 (51.5%) | 12,377 (51.6%) |
| Male | 11,489 (48.5%) | 11,616 (48.4%) |
| Ethnicity, *n* (%) |  |  |
| Chinese | 20,391 (86.2%) | 20,432 (85.2%) |
| Indian | 1,177 (5.0%) | 1,349 (5.6%) |
| Malay | 737 (3.1%) | 778 (3.2%) |
| Others | 1,363 (5.8%) | 1,434 (6.0%) |
| Citizenship, *n* (%) |  |  |
| Singaporean / Permanent Resident | 21,396 (90.4%) | 21,645 (90.2%) |
| Others | 2,272 (9.6%) | 2,348 (9.8%) |
| Class year, *n* (%) |  |  |
| 1 | 7,359 (31.1%) | 7,457 (31.1%) |
| 2 | 6,165 (26.0%) | 6,897 (28.7%) |
| 3 | 4,593 (19.4%) | 4,561 (19.0%) |
| 4 | 5,160 (21.8%) | 4,812 (20.1%) |
| >4 | 391 (1.7%) | 266 (1.1%) |
| **Class information** |  |  |
| No. of course modules offered | 1,816 | 1,734 |
| No. of class/tutorial groups | 6,104 | 5,929 |
| **Wi-Fi connection information** |  |  |
| Students who used Wi-Fi, *n* (%) | 23,586 (99.7%) | 23,881 (99.5%) |
| No. of Wi-Fi access points connected | 6,573 | 6,313 |
| No. of Wi-Fi connections | 19,282,203 | 4,871,285 |

**Extended Data Table 2.**
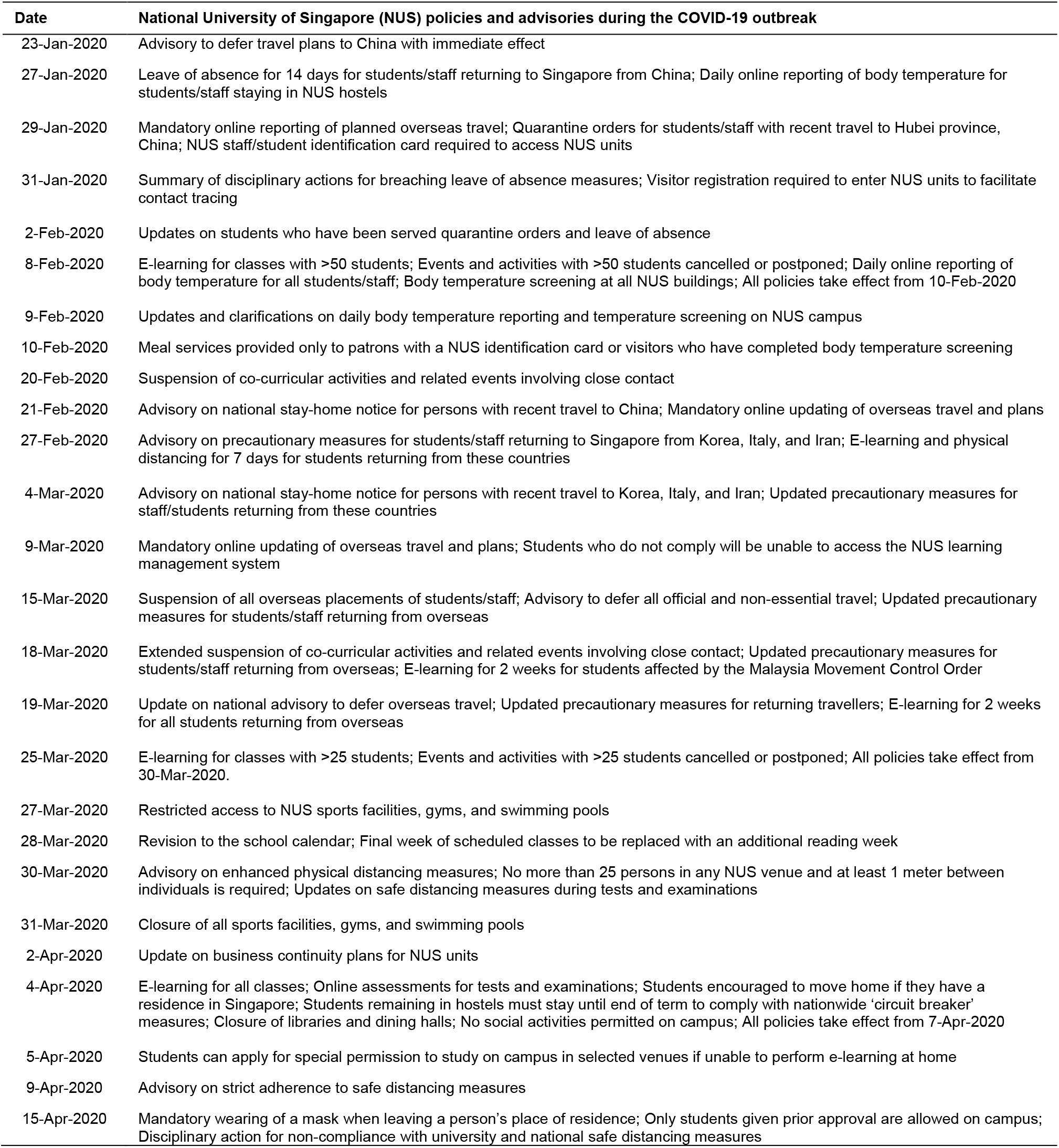
University policies and advisories during the COVID-19 outbreak.

| Date | National University of Singapore (NUS) policies and advisories during the COVID-19 outbreak |
| --- | --- |
| 23-Jan-2020 | Advisory to defer travel plans to China with immediate effect |
| 27-Jan-2020 | Leave of absence for 14 days for students/staff returning to Singapore from China; Daily online reporting of body temperature for students/staff staying in NUS hostels |
| 29-Jan-2020 | Mandatory online reporting of planned overseas travel; Quarantine orders for students/staff with recent travel to Hubei province, China; NUS staff/student identification card required to access NUS units |
| 31-Jan-2020 | Summary of disciplinary actions for breaching leave of absence measures; Visitor registration required to enter NUS units to facilitate contact tracing |
| 2-Feb-2020 | Updates on students who have been served quarantine orders and leave of absence |
| 8-Feb-2020 | E-learning for classes with >50 students; Events and activities with >50 students cancelled or postponed; Daily online reporting of body temperature for all students/staff; Body temperature screening at all NUS buildings; All policies take effect from 10-Feb-2020 |
| 9-Feb-2020 | Updates and clarifications on daily body temperature reporting and temperature screening on NUS campus |
| 10-Feb-2020 | Meal services provided only to patrons with a NUS identification card or visitors who have completed body temperature screening |
| 20-Feb-2020 | Suspension of co-curricular activities and related events involving close contact |
| 21-Feb-2020 | Advisory on national stay-home notice for persons with recent travel to China; Mandatory online updating of overseas travel and plans |
| 27-Feb-2020 | Advisory on precautionary measures for students/staff returning to Singapore from Korea, Italy, and Iran; E-learning and physical distancing for 7 days for students returning from these countries |
| 4-Mar-2020 | Advisory on national stay-home notice for persons with recent travel to Korea, Italy, and Iran; Updated precautionary measures for staff/students returning from these countries |
| 9-Mar-2020 | Mandatory online updating of overseas travel and plans; Students who do not comply will be unable to access the NUS learning management system |
| 15-Mar-2020 | Suspension of all overseas placements of students/staff; Advisory to defer all official and non-essential travel; Updated precautionary measures for students/staff returning from overseas |
| 18-Mar-2020 | Extended suspension of co-curricular activities and related events involving close contact; Updated precautionary measures for students/staff returning from overseas; E-learning for 2 weeks for students affected by the Malaysia Movement Control Order |
| 19-Mar-2020 | Update on national advisory to defer overseas travel; Updated precautionary measures for returning travellers; E-learning for 2 weeks for all students returning from overseas |
| 25-Mar-2020 | E-learning for classes with >25 students; Events and activities with >25 students cancelled or postponed; All policies take effect from 30-Mar-2020. |
| 27-Mar-2020 | Restricted access to NUS sports facilities, gyms, and swimming pools |
| 28-Mar-2020 | Revision to the school calendar; Final week of scheduled classes to be replaced with an additional reading week |
| 30-Mar-2020 | Advisory on enhanced physical distancing measures; No more than 25 persons in any NUS venue and at least 1 meter between individuals is required; Updates on safe distancing measures during tests and examinations |
| 31-Mar-2020 | Closure of all sports facilities, gyms, and swimming pools |
| 2-Apr-2020 | Update on business continuity plans for NUS units |
| 4-Apr-2020 | E-learning for all classes; Online assessments for tests and examinations; Students encouraged to move home if they have a residence in Singapore; Students remaining in hostels must stay until end of term to comply with nationwide 'circuit breaker' measures; Closure of libraries and dining halls; No social activities permitted on campus; All policies take effect from 7-Apr-2020 |
| 5-Apr-2020 | Students can apply for special permission to study on campus in selected venues if unable to perform e-learning at home |
| 9-Apr-2020 | Advisory on strict adherence to safe distancing measures |
| 15-Apr-2020 | Mandatory wearing of a mask when leaving a person's place of residence; Only students given prior approval are allowed on campus; Disciplinary action for non-compliance with university and national safe distancing measures |

**Extended Data Fig. 1.**
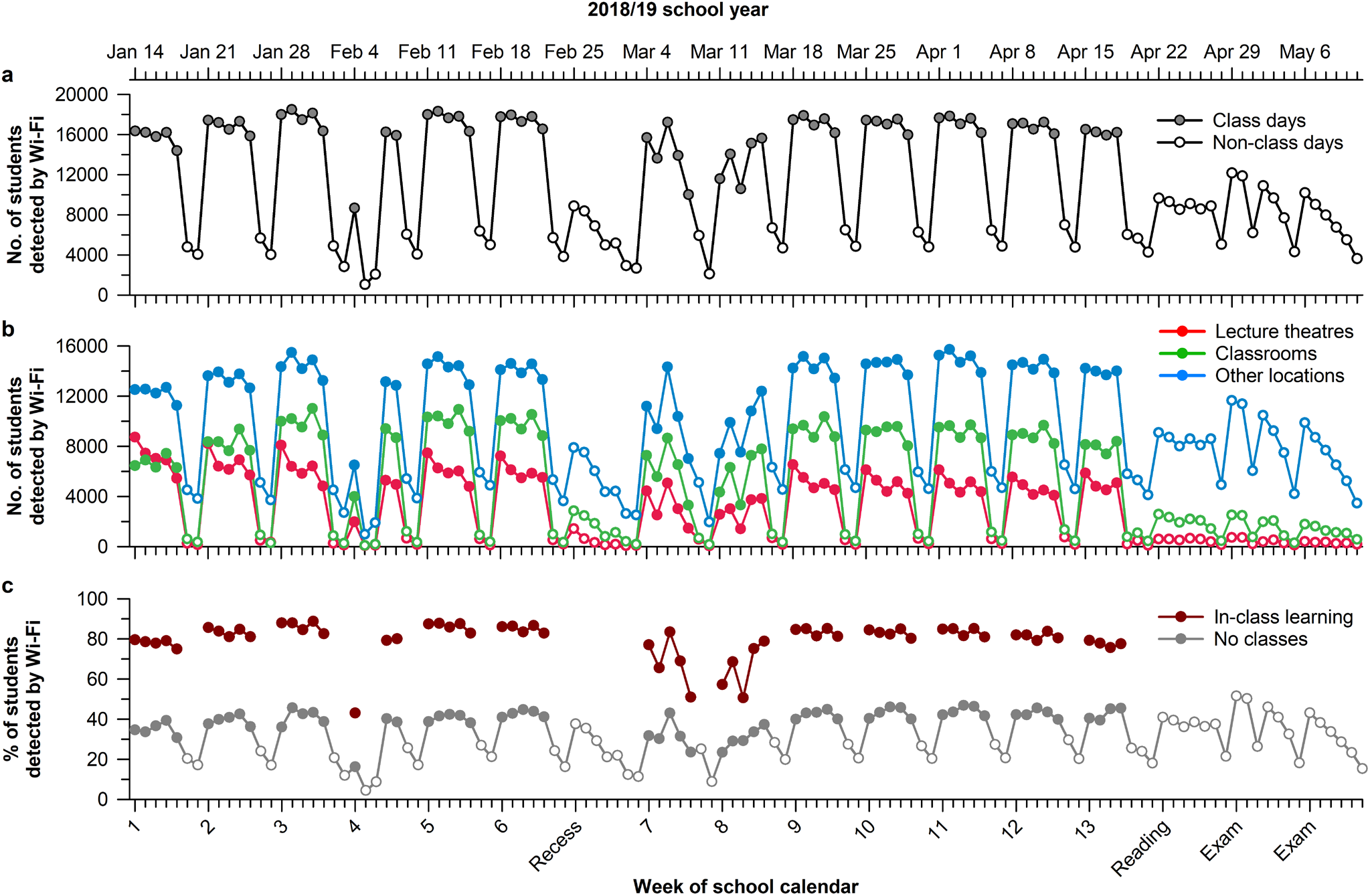
Students detected on campus during normal university operations. Data are shown for the second semester of the 2018/19 school year at the National University of Singapore (NUS), assessed one year before the COVID-19 outbreak. The number of students per day who connected to the NUS Wi-Fi network is shown for (**a**) the entire campus and (**b**) different types of locations on campus. (**c**) The daily percentage of students detected by Wi-Fi was about two-fold greater in students with in-class learning versus no scheduled class. In panels **b** and **c**, open circles indicate non-class days.

**Extended Data Fig. 2.**
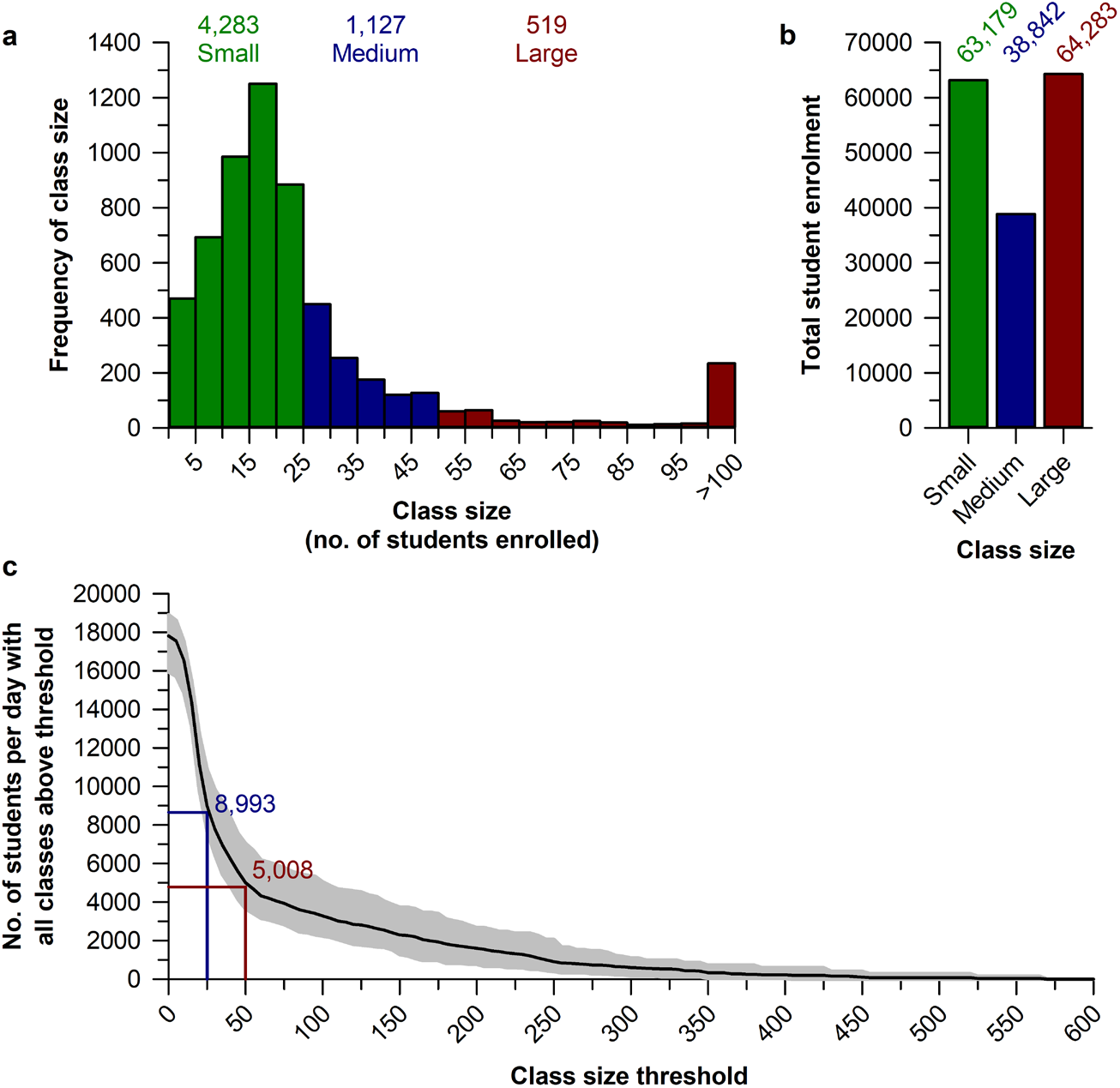
Class size characteristics at the National University of Singapore (NUS). (**a**) The distribution of class sizes is shown for the second semester of the 2019/20 school year in which the COVID-19 outbreak occurred. Class sizes were categorized as small (green; ≤25 students), medium (blue; >25 to ≤50 students), or large (red; >50 students). (**b**) The combined student enrolment in medium and large classes was greater than enrolment in small classes. (**c**) The cumulative distribution plot shows the number of students whose smallest class of the day exceeded a given class size threshold. The black trace with shaded grey lines shows the daily mean and range. The red dropline shows that the transition to e-learning for classes with >50 students resulted in about 5,000 students per day who had classes delivered only by e-learning. The blue dropline shows that the transition to e-learning for classes with >25 students resulted in about 9,000 students per day who had classes delivered only by e-learning. When all classes were shifted to e-learning there were about 18,000 students per day taking their classes online.

**Extended Data Fig. 3.**
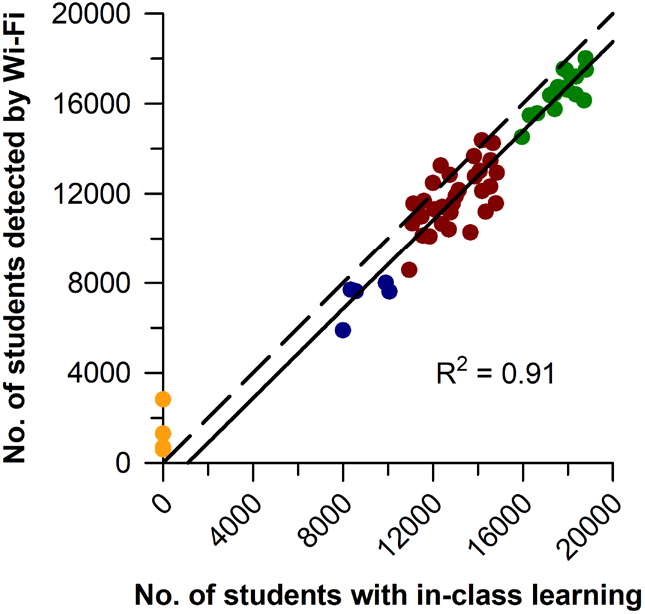
The daily number of students detected on campus was predicted by the number of students with in-class learning. Data are shown for the second semester of the 2019/20 school year at the National University of Singapore (NUS) during the COVID-19 outbreak. The number of students per day who connected to the NUS Wi-Fi network is plotted against the daily number of students who had at least one class session that took place on campus. Circle colours correspond to different parts of the semester with normal in-class learning (green), e-learning for classes with >50 students (red), e-learning for classes with >25 students (blue), and e-learning for all classes (orange). The solid black trace shows the best-fit linear regression model, and the dashed black trace is the unity line.

**Extended Data Fig. 4.**
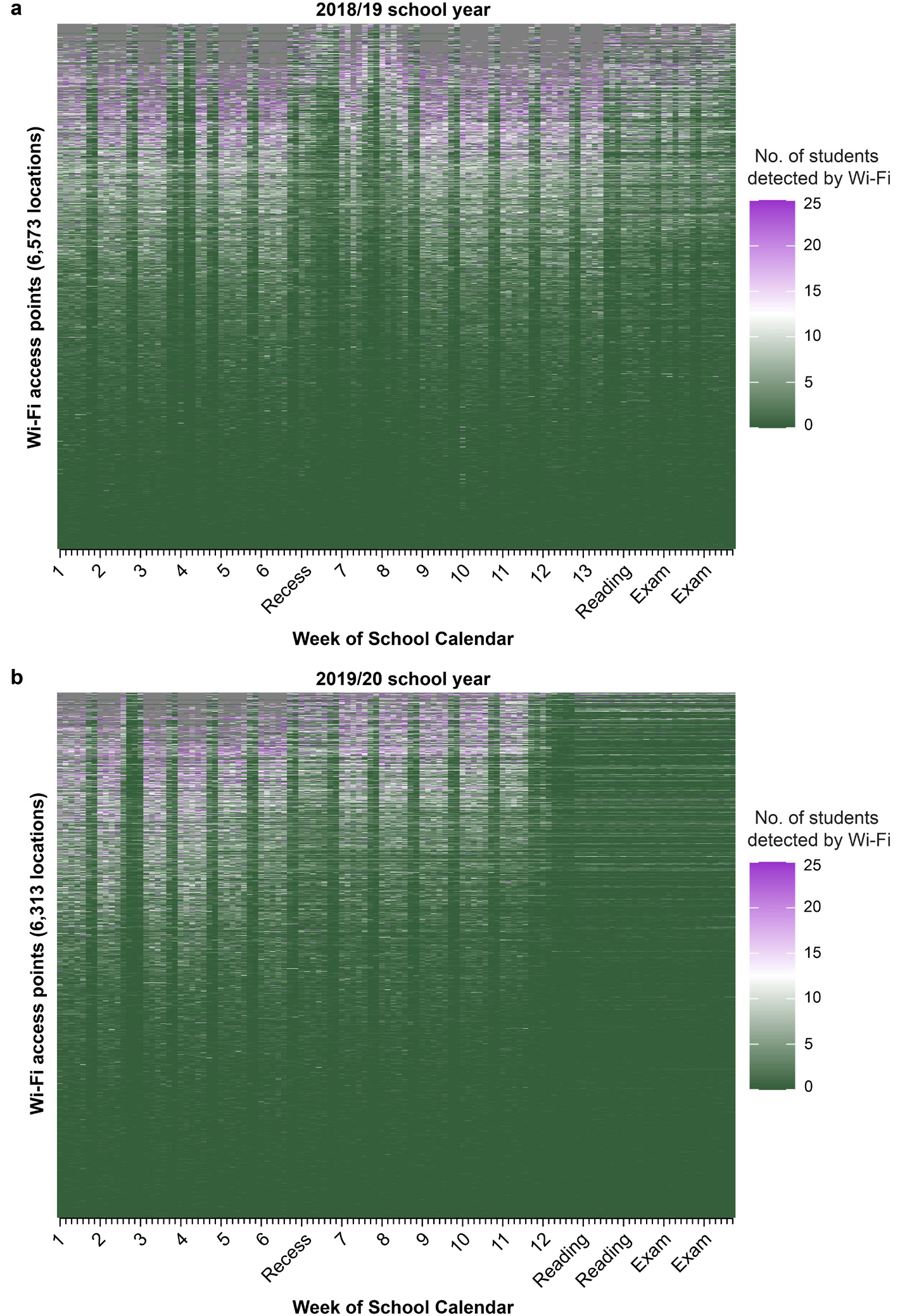
Campus-wide detection of students at different Wi-Fi access points at the National University of Singapore (NUS). The daily peak in the number of students who connected to each Wi-Fi access point is shown for (**a**) the second semester of the 2018/19 school year, and (**b**) the second semester of the 2019/20 school year in which the COVID-19 outbreak occurred. Each peak value corresponds to largest number of students per day detected at a given Wi-Fi access point over a 15-min period. Each row in the heat map represents a different Wi-Fi access point with green and magenta colours indicating the number of students who were detected.

**Extended Data Fig. 5.**
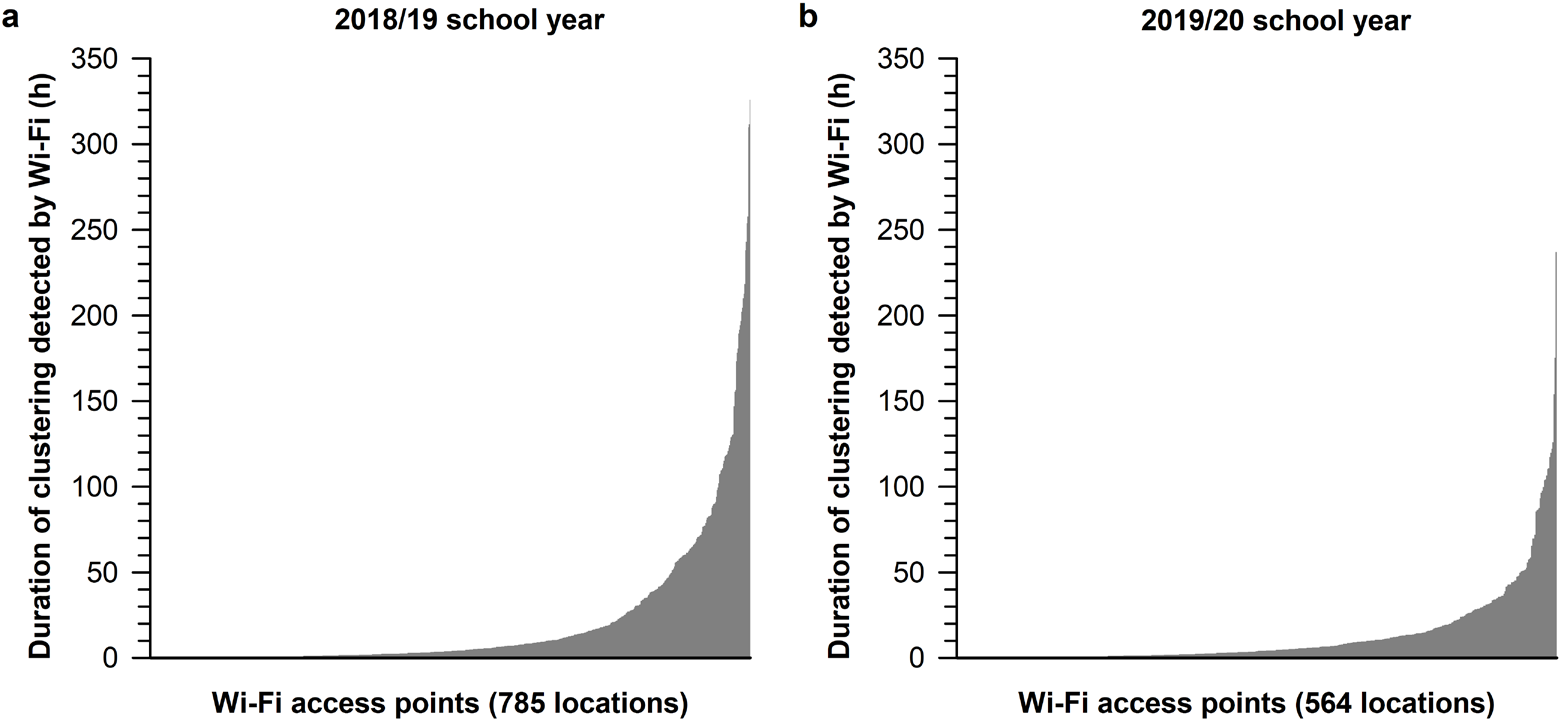
Distribution of student clustering across Wi-Fi access points at the National University of Singapore (NUS). The cumulative duration of student clustering (>25 students connected to the same Wi-Fi access point) is shown for (**a**) the second semester of the 2018/19 school year, and (**b**) the second semester of the 2019/20 school year in which the COVID-19 outbreak occurred. Data are plotted for Wi-Fi access points with at least one student cluster detected during the semester (785 out of 6,573 locations in 2018/19; 564 out of 6,313 locations in 2019/20). Wi-Fi access points in each plot are ordered from left to right by the cumulative duration of student clustering over the entire semester.

**Extended Data Fig. 6.**
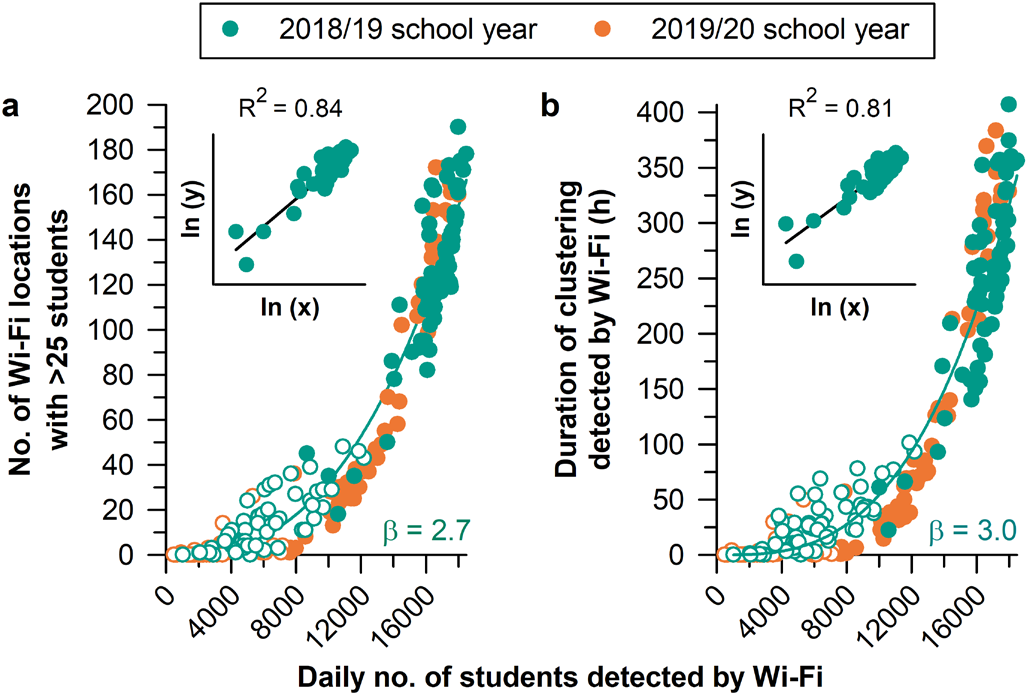
Student clustering showed power law scaling with the number of students on campus. Students’ Wi-Fi connection data were analysed for the second semester of the 2018/19 school year and compared with the second semester of the 2019/20 school year in which the COVID-19 outbreak occurred. In both semesters, student clustering behaviour showed accelerated growth with increasing number of students detected on campus, including (**a**) the number of Wi-Fi locations with a student cluster (>25 students connected to the same Wi-Fi access point), and (**b**) the duration of student clustering at these locations. Each dataset was fitted with a power law function, with β representing the scaling exponent. Insets show results for linear regression after taking the natural logarithm of each variable for the 2018/19 school year. Filled circles show school days and open circles indicate non-class days.

**Extended Data Fig. 7.**
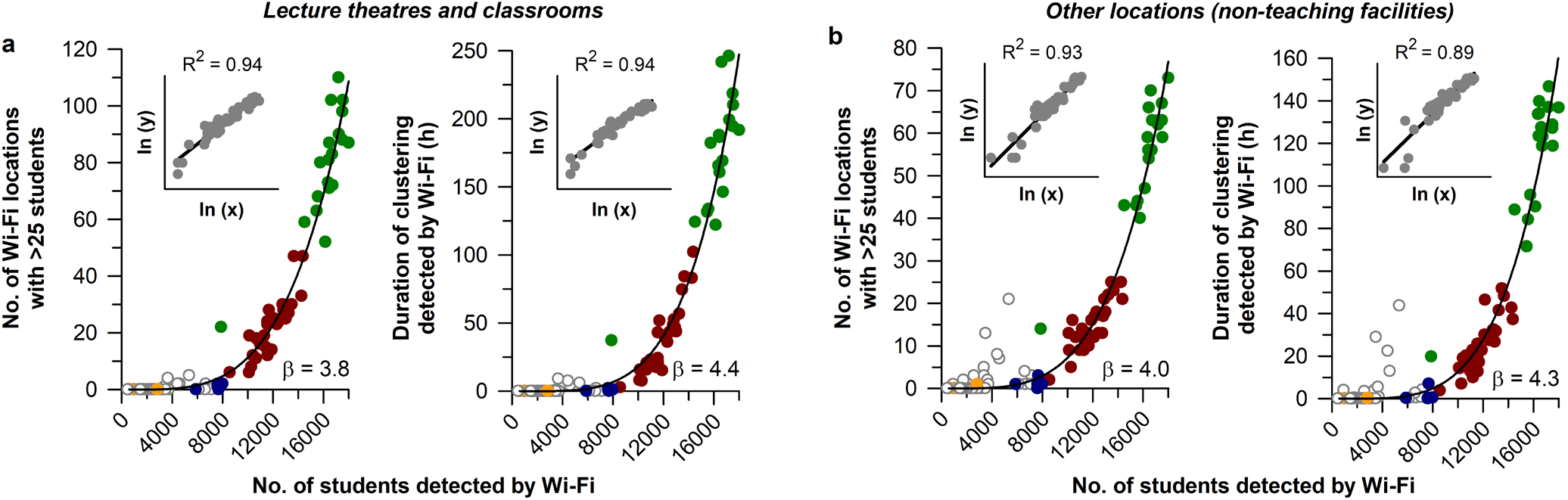
Student clustering at different campus locations showed power law scaling with the number of students detected on campus. Data are shown for the second semester of the 2019/20 school year at the National University of Singapore (NUS) during the COVID-19 outbreak. In both (**a**) teaching facilities and (**b**) non-teaching facilities, the number of Wi-Fi locations with >25 students (left panels) and the duration of clustering behaviour (right panels) showed accelerated growth with increasing number of students detected on campus. Each dataset was fitted with a power law function, with β representing the scaling exponent. Insets show results for linear regression after taking the natural logarithm of each variable. Circle colours correspond to different parts of the semester with normal in-class learning (green), e-learning for classes with >50 students (red), e-learning for classes with >25 students (blue), and e-learning for all classes (orange). Open circles indicate non-class days.

**Extended Data Fig. 8.**
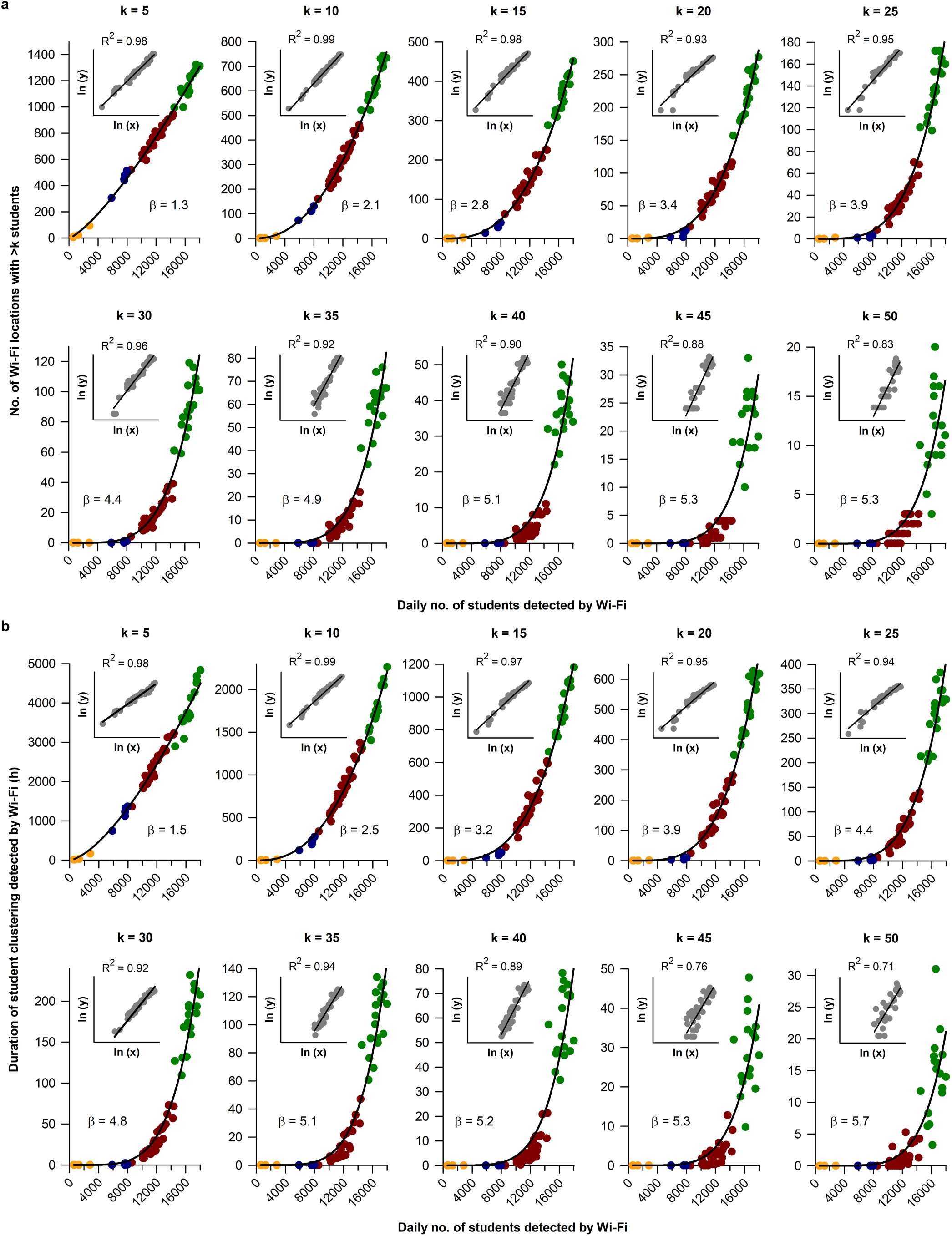
Power law scaling of different student cluster sizes with number of students detected on campus. Data are shown for the second semester of the 2019/20 school year at the National University of Singapore (NUS) during the COVID-19 outbreak. Different definitions of a student cluster were tested ranging from >5 to >50 students detected at the same Wi-Fi access point. For all cluster sizes, student clustering behaviour showed accelerated growth with increasing number of students detected on campus, including (**a**) the number of Wi-Fi locations with a student cluster, and (**b**) the duration of student clustering at these locations. Each dataset was fitted with a power law function, with β representing the scaling exponent. Insets show results for linear regression after taking the natural logarithm of each variable. Circle colours correspond to different parts of the semester with normal in-class learning (green), e-learning for classes with >50 students (red), e-learning for classes with >25 students (blue), and e-learning for all classes (orange). Open circles indicate non-class days.

